# Alternative end-joining results in smaller deletions in heterochromatin relative to euchromatin

**DOI:** 10.1101/2023.03.03.531058

**Authors:** Jacob M. Miller, Sydney Prange, Huanding Ji, Alesandra R. Rau, Varandt Y. Khodaverdian, Xiao Li, Avi Patel, Nadejda Butova, Avery Lutter, Helen Chung, Chiara Merigliano, Chetan C. Rawal, Terrence Hanscom, Mitch McVey, Irene Chiolo

**Author notes:** Equal contribution (Jacob M. Miller, Sydney Prange, Huanding Ji, Alesandra R. Rau).

## Abstract

Pericentromeric heterochromatin is highly enriched for repetitive sequences prone to aberrant recombination. Previous studies showed that homologous recombination (HR) repair is uniquely regulated in this domain to enable ‘safe’ repair while preventing aberrant recombination. In *Drosophila* cells, DNA double-strand breaks (DSBs) relocalize to the nuclear periphery through nuclear actin-driven directed motions before recruiting the strand invasion protein Rad51 and completing HR repair. End-joining (EJ) repair also occurs with high frequency in heterochromatin of fly tissues, but how alternative EJ (alt-EJ) pathways operate in heterochromatin remains largely uncharacterized. Here, we induce DSBs in single euchromatic and heterochromatic sites using a new system that combines the DR-*white* reporter and I-SceI expression in spermatogonia of flies. Using this approach, we detect higher frequency of HR repair in heterochromatin, relative to euchromatin. Further, sequencing of mutagenic repair junctions reveals the preferential use of different EJ pathways across distinct euchromatic and heterochromatic sites. Interestingly, synthesis-dependent microhomology-mediated end joining (SD-MMEJ) appears differentially regulated in the two domains, with a preferential use of motifs close to the cut site in heterochromatin relative to euchromatin, resulting in smaller deletions. Together, these studies establish a new approach to study repair outcomes in fly tissues, and support the conclusion that heterochromatin uses more HR and less mutagenic EJ repair relative to euchromatin.

## INTRODUCTION

Pericentromeric heterochromatin (hereby ‘heterochromatin’) occupies ∼6-90% of eukaryotic genomes, or ∼30% of the fly genome, and is mostly composed of highly repeated DNA sequences [1-6]. In *Drosophila* about half of these sequences are satellite repeats (mostly 5-bp sequences repeated in tandem for hundreds of kilobases to megabases), and the rest is transposons, other scrambled repeats, and only about 250 genes [1, 3]. Given the abundance of repeated sequences, heterochromatin is prone to aberrant recombination during DSB repair (reviewed in [7]).

DSBs can be repaired through different pathways, mostly: HR, single-strand annealing (SSA), classical non-homologous end joining (cNHEJ), and alternative-EJ (alt-EJ) (reviewed in [8]). During HR, DSBs are resected to generate long stretches of single-stranded DNA (ssDNA) that invade homologous sequences on the sister chromatid or the homologous chromosome, which are used as templates for DNA synthesis and repair. HR is typically error-free in single copy sequences. In heterochromatin, however, the presence of thousands to millions of alternative donor sequences associated with different chromosomes can lead to strand invasion of non-allelic (ectopic) templates, resulting in aberrant recombination, chromosome rearrangements and genome instability.

Previous studies in *Drosophila* cultured cells showed that HR repair of heterochromatic DSBs occurs through a dedicated pathway that relocalizes repair sites to the nuclear periphery. Resection starts inside the heterochromatin domain [9], but the strand invasion protein Rad51 is recruited only after relocalization [9-11]. This movement occurs by directed motion along a network of nuclear actin filaments (F-actin), and is mediated by nuclear myosins associated with the Smc5/6 complex at repair sites [12-15]. F-actin-dependent relocalization of heterochromatic DSBs to outside the ‘chromocenters’ has also been detected in mouse cells [12, 16, 17], revealing conserved pathways. This mechanism likely promotes ‘safe’ repair by moving the damage away from the bulk of ectopic repeated sequences inside the heterochromatin domain, before promoting strand invasion with the homologous templates on the sister chromatid or the homologous chromosome (reviewed in [18, 19]). In flies, homologous pairing occurs throughout interphase, thus the homologous chromosome is readily available as a template for HR repair [20]. Accordingly, both the sister chromatid and the homologous chromosome can be used as repair templates for HR in heterochromatin, although the sister chromatid is used at higher frequency [21]. Studies in fly and human cultured cells suggest that HR is the preferred pathway used for heterochromatin repair when both HR and NHEJ are available, *e.g.,* in S and G2 phases of the cell cycle [9, 22].

In addition to HR, long-range resection can promote SSA when ssDNA unmasks identical sequences that can pair with each other, resulting in large deletions (reviewed in [23, 24]). Despite the abundance of repeated sequences in heterochromatin, SSA seems rarely used in this domain [21].

Alternatively, short-range resection and small regions of homology (1-8bp) can be used in pol theta (polθ)-dependent alt-EJ pathways: microhomology-mediated EJ (MMEJ) or synthesis-dependent MMEJ (SD-MMEJ) [25-33] (reviewed in [34-36]). In MMEJ, the annealing of regions of homology available on the ssDNA after resection, is followed by flap removal, fill-in synthesis and ligation, resulting in small deletions [26]. Conversely, SD-MMEJ is characterized by annealing followed by a synthesis step, dissociation, and a re-annealing using the nascent DNA before fill-in synthesis and ligation, frequently resulting in indels (deletions with insertions) [25, 35, 37]. Finally, cNHEJ simply rejoins the two ends of the DSB together through binding of Ku70/Ku80, processing by nucleases and polymerases, and ligation by a complex containing DNA Ligase 4 (Lig4) (reviewed in [38]). cNHEJ can be error-free or associated with small alterations at the junction (reviewed in [38-40]), with error-free outcomes prevailing in a context of ‘clean’ breaks (*e.g.*, those induced by endonucleases) (reviewed in [41]). Given the different mutagenic outcomes of these repair pathways, it is possible to detect their activity based on the sequence of the repair product [42-44].

A major determinant of EJ outcomes is the DNA sequence surrounding the DSB [37, 42, 45] (reviewed in [40]). Alt-EJ is promoted by the presence of small regions of homology around the cut site, with SD-MMEJ particularly facilitated by the establishment of secondary structures that promote the first synthesis step [37, 42, 45]. In agreement, the binding of replication protein A (RPA) antagonizes alt-EJ by preventing ssDNA annealing and secondary structure formation [46, 47]. While much is known about the role of the DNA sequence in alt-EJ pathways, the contribution of the chromatin context remains poorly understood.

A recent study systematically looked at the balance between MMEJ and cNHEJ in different epigenetic environments using a sequencing-based reporter cut with Cas9 inserted in a large number of genomic locations of human cells [43]. This study identified a higher ratio of MMEJ:cNHEJ usage in H3K9me2-enriched lamina-associated silenced regions, and in polycomb silenced regions (H3K27me3-enriched), relative to active chromatin [43]. This appears to result from more efficient usage of cNHEJ in active chromatin, suggesting that silencing interferes with cNHEJ while enabling MMEJ [43]. Thus, this study revealed that epigenetic environments can influence EJ repair outcomes, but it did not examine SD-MMEJ repair, nor did it investigate DSB repair in pericentromeric heterochromatin. This nuclear domain is unique in that it is not only enriched for H3K9me3 [2, 9, 48], but is also mostly composed of repeated sequences [1, 3], is phase separated [49, 50], and is not typically associated with the nuclear periphery (reviewed in [5, 19, 51]).

A major advantage of using the same reporter inserted in different genomic locations is that repair outcomes are not affected by differences in the local DNA sequence, thus enabling a more direct focus on effects associated with chromatin composition and nuclear domains. Previous studies established the DR-*white* reporter as an excellent tool to study both HR and EJ pathways in *Drosophila* [21, 52]. *Drosophila* is also an ideal model system to study alt-EJ, given that this pathway is commonly used to repair DSBs even in the presence of a fully functional cNHEJ [44, 53-55].

Here, we use a DR-*white* reporter inserted in euchromatic and heterochromatic locations of *Drosophila*, and we directly investigate repair outcomes in spermatogonia and primary spermatocytes as a model for mitotically dividing cells of adult flies. Contrary to previous conclusions, we detect higher frequency of HR repair in all heterochromatic insertions relative to euchromatin. In addition, different alt-EJ repair mechanisms prevail at distinct heterochromatic locations, with SD-MMEJ typically resulting in smaller deletions in heterochromatin relative to euchromatin. This is linked to a higher usage of repeat motifs closer to the cut site, suggesting that farther motifs are less accessible for secondary structure formation in heterochromatin and/ or that repair protein access is different in the two chromatin contexts. Together, this study establishes new tools to study heterochromatin repair in flies, reveals unique features of alt-EJ in heterochromatin, and suggests that alt-EJ is less mutagenic in this domain. Most surprisingly, this study supports a model where the chromatin context, not just the sequence composition, affects motif usage for SD-MMEJ repair.

## RESULTS

### A new approach to induce single DSBs in euchromatin or heterochromatin of spermatogonia

We generated new fly lines to examine repair outcomes with the DR-*white* reporter. This reporter is characterized by two non-functional repeats of the *white* gene (*Sce.white* and *iwhite*), the first of which contains a restriction site for I-SceI (Fig. 1A) [52]. Exposure to the I-SceI endonuclease results in DSB induction in *Sce.white*. Intra-chromosomal or inter-sister chromatid HR repair using the *iwhite* sequence as a template restores the *wild-type white* gene. SSA repair results in the loss of the DsRed fluorescent marker. Mutagenic EJ repair results in the loss of the I-SceI cut site and can be characterized through PCR amplification across the cut site, enzymatic digestion, and DNA sequencing [52].

**Figure 1:**
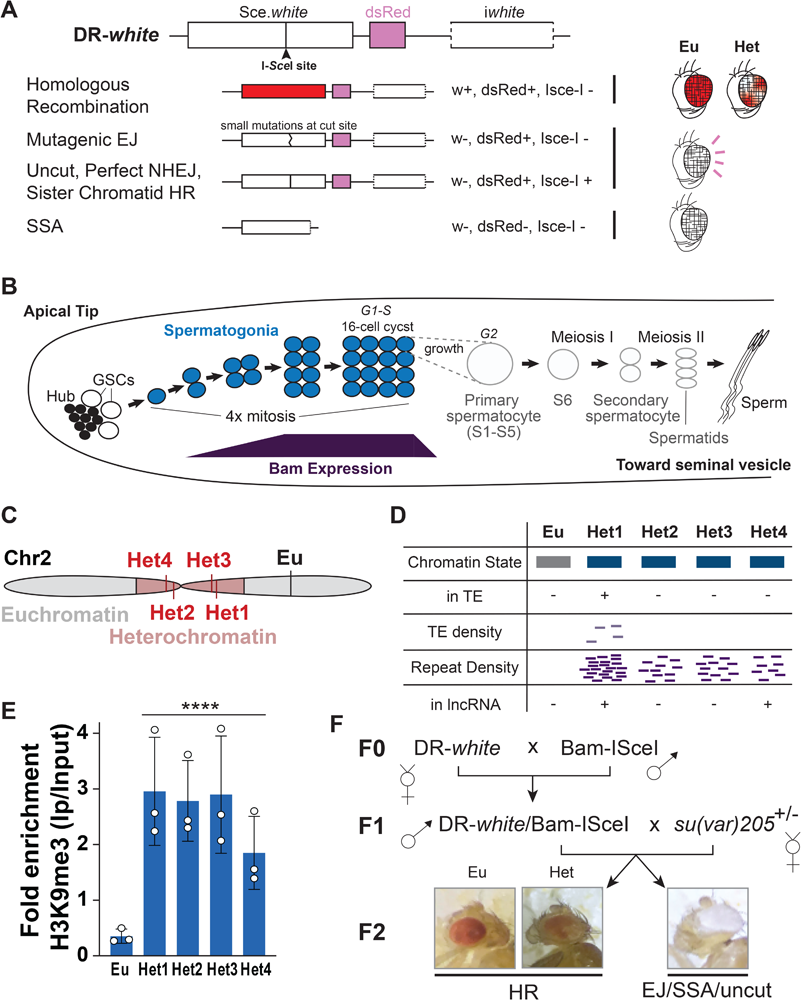
Experimental setup used to induce single DSBs in spermatogonia. A) Schematic representation of the DR-*white* reporter, containing a *white* gene with an inserted I-SceI site (*Sce.white*) that introduces a premature stop codon, the dsRed sequence, and a truncated *white* gene (*iwhite*) [52]. dsRed and *white* are driven by the eye tissue-specific 3xP3 promoter and glass multiple reporter (GMR), respectively. HR repair of I-SceI induced breaks using the *iwhite* sequence in the germline, results in *w+*, *i.e.,* flies with red eyes (for *DR-white* insertions in euchromatin) or variegated-red eyes (for *DR-white* insertions in heterochromatin) in the progeny. SSA repair results in the loss of the fluorescent marker dsRed in the eyes. Mutagenic repair, absence of cut or perfect NHEJ results in white-eyed flies (*w-*), which can be further characterized by PCR amplification, I-SceI digestion, and sequencing across the repair junction. B) Schematic representation of the testis, highlighting the spermatogonia and the corresponding level of activity of the *Bam* promoter. C) Schematic representation of the genomic location of the DR-*white* insertions used in this study. D) Description of some of the most distinctive features of these sites. Colors indicate the chromatin state described by the 9-state model [61]. Blue: Pericentromeric heterochromatin. Grey: ‘background’ state, typically corresponding to large domains between active genes. Het1 is located in a transposable element (TE). TE density and repeat density are qualitatively illustrated by the number of light and dark purple bars, respectively. 1 bar: 2 TEs or 10 repeats in a 20Kb window around the insertion site. Het1 and Het4 insertion sites are also located inside of an annotated lncRNA (+), while Eu, Het3, and Het2 are not (-). E) H3K9me3 levels at the insertion sites were determined by ChIP-seq data available for Kc cells. Enrichments are calculated as the ratio of reads +/- 0.5 Mb from the insertion site per million mapped reads between IP and input samples. ****p<0.0001 by two-sample z-test for means. Error bars: SD of 3 replicates. F) Scheme of the fly crosses set up to analyze repair outcomes in the F2 generation.

We induced I-SceI expression using the *Bam* promoter (BamP898-CFI [56]), which is activated at the 2-cell cyst spermatogonia stage, peaks at the 8 cell-cyst stage, and is quickly suppressed in spermatocytes [57, 58] (Fig. 1B). In this transgenic construct, I-SceI protein levels are also suppressed at the translational level in spermatocytes through the Bam 3’ UTR. This confines I-SceI expression and DSB induction to male pre-meiotic germ cells. This is an important advancement from previous studies that expressed I-SceI broadly in the germ line or in somatic cells using a heat-shock (hs) or a ubiquitin (Ub) promoter [21, 52, 59]. Targeted expression in spermatogonia [57, 58], enables damage induction and repair in a homogeneous population of mitotically dividing cells, reducing confounding effects from multiple cell types and stages of differentiation. In addition, inducing I-SceI via hs or Ub promoters that are active in the germinal stem cells can give rise to ‘jackpot’ events that populate the entire germ line, introducing significant bias in the frequency of repair outcomes [52, 60]. These events are not induced by the *Bam* promoter, eliminating another major source of ‘noise’ in the system. With this approach, each individual of the progeny contains a unique repair product that can be scored for the acquisition of *white+* (red eyes) or DsRed-(red fluorescent eyes) phenotypes and further analyzed by sequencing.

We integrated DR-*white* in one euchromatic location (Eu) and four pericentromeric heterochromatic sites (Het1-4) of chromosome 2 (Fig. 1C, D). The euchromatic site was previously studied with the DR-*white* cassette and its genomic location is characterized by a ‘gray’ chromatin state (or ‘state 9’) in the 9-state model of combinatorial chromatin patterns [61], which corresponds to regions deprived of active or silent histone marks and which are typically interspersed with active genes (Fig. 1D). Consistently, very low levels of H3K9me3 were detected at this site (Fig. 1E) [62, 63]. The *yellow*+ marker of the MiMIC line used to generate this insertion and the Ds-Red and *white+* marker of the DR-*white* reporter are fully expressed under the respective promoters, in agreement with the lack of silencing effects. Conversely, the heterochromatic sites are heavily silenced based on H3K9me3 levels (Fig. 1E), and are characterized by a ‘blue’ chromatin state [61] (Fig. 1D), corresponding to silenced pericentromeric heterochromatin. These locations are also characterized by a high number of repeats surrounding the insertion site (Fig. 1D). Importantly, the yellow marker of the MiMIC lines used to generate these insertions is nearly fully silenced resulting in a faint ‘mosaic’ phenotype, while the Ds-Red and the *white+* markers of the DR-*white* reporter are completely silenced. This indicates that silent histone marks spread across the cassette, creating a silenced epigenetic context at the cut site.

Given the silent state of the heterochromatic insertions, F1 males were crossed with females carrying a mutation in the heterochromatin silencing component HP1a (*su(var)205+/-*) [64] to unmask the presence of the reporter (Fig. 1F). HP1 heterozygosity in the F2 generation results in partial reactivation of DsRed and *white* markers carried by the reporter (Fig. 1F), leading to a mosaic phenotype that can easily be scored to assess repair outcomes. Thus, with this system, DSB induction and repair occur in the spermatogonia of F1 males, and can be scored in the F2 progeny.

### HR repair occurs more frequently in heterochromatin

First, we investigated HR repair of adult F1 males containing DR-*white* and *Bam*-ISceI by quantifying the fraction of individuals with red eyes (w+, DsRed+) in the progeny (Fig. 1F).

This analysis revealed that 3.6% (+/- 0.3%) of the flies repair DSBs using HR when DR-*white* is placed in euchromatin. This percentage remains remarkably consistent across replicates, in agreement with little experimental variability. This is a significantly lower frequency than observed in previous studies with I-SceI under a heat-shock promoter and DR-*white* in the same genomic location, where HR occurred at a frequency of ∼20% [21]. The lower HR frequency is consistent with *Bam* being active only in a subset of cells in the germline and thus inducing DSBs less frequently.

Next, we analyzed HR repair in the heterochromatic insertions of DR-*white* by scoring variegated red-eyed flies in the F2 generation. All heterochromatic insertions used HR at significantly higher frequency relative to the euchromatic insertion, ranging from 5.5% to 9.2% of the total DR-*white*-containing F2 progeny (Fig. 2). This corresponds to ∼1.5-2.5 fold increase in HR frequency for heterochromatic DR-*white* insertions relative to the euchromatic insertion. Thus, we conclude that HR repair is more frequently used in heterochromatin relative to euchromatin.

**Figure 2:**
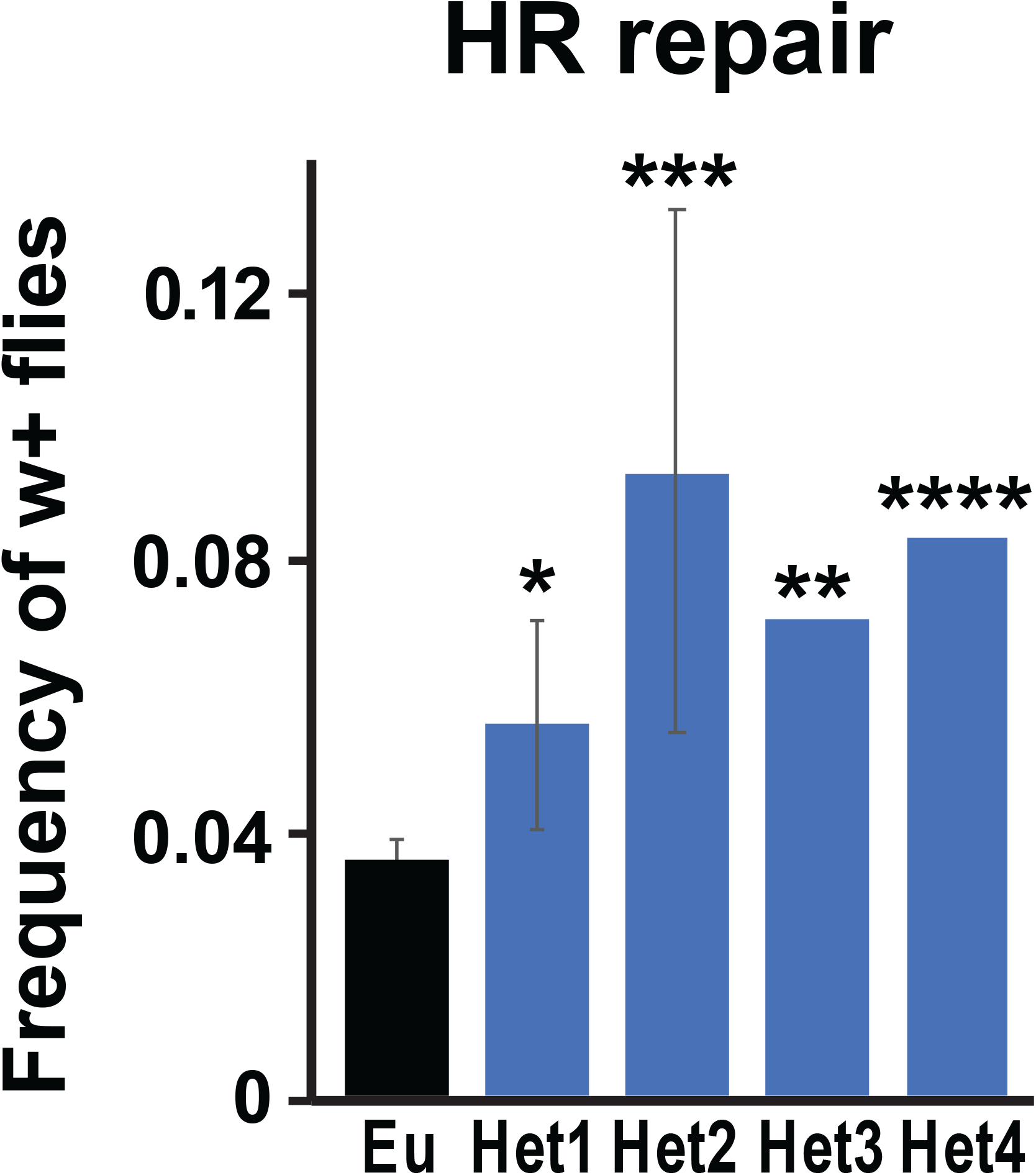
Frequency of HR products. The frequency of HR products for indicated DR-*white* insertions was calculated as the number of red-eyed flies over the total number of flies (n) in the F2 progeny. *p=0.015; **p=0.007; ***p=0.0002; ****p<0.0001 by one-tailed unpaired t-test. Flies analyzed: n=1352 for Eu; n=1267 for Het1; n=880 for Het2; n=679 for Het3; n=485 for Het4. Error bars: SD of two independent experiments when available.

### Alt-EJ repair pathways operate with different frequencies at distinct DR-white insertions

We then focused on characterizing EJ events by analyzing repair products in white-eyed flies. White-eyed F2 individuals can derive from SSA (w-DsRed-) or mutagenic EJ repair (w-DsRed+), resulting in loss of the I-SceI cut site (Fig. 1A). Alternatively, white-eyed flies can result from germ line cells where the I-SceI site was not cut, from perfect NHEJ repair, or from HR with the sister chromatid, and each of these outcomes retain a wild-type I-SceI site (Fig. 1A). Mutagenic and non-mutagenic events were distinguished through PCR amplification across the cut site followed by digestion with I-SceI (Fig. 1A).

Consistent with previous studies [21, 52, 59], this analysis showed that SSA is rarely used in euchromatin and heterochromatin (Suppl. Fig. 1A) and its frequency is not significantly different across DR-*white* insertions. Conversely, mutagenic EJ repair occurs with high frequency across all DR-*white* insertions, accounting for ∼20-50% of all the events (Suppl. Fig. 1B).

We also identified a large number of flies where the I-SceI site was still intact (∼40-60% of the white-eyed flies, Suppl. Fig. 1B), consistent with either low cut efficiency or non-mutagenic repair. The frequency of mutagenic and non-mutagenic events is similar across euchromatic and heterochromatic DR-*white* insertions, with the exception of Het4 where non-mutagenic outcomes are more frequent potentially reflecting lower cut efficiency at this site (Suppl. Fig. 1B). Of note, observing higher frequency of EJ relative to HR is consistent with other studies with the DR-*white* cassette [21, 52]. This likely reflects the inability of the DR-*white* to detect intersister HR resulting in perfect repair, which is the preferred mechanism for HR repair [8, 65].

We further characterized mutagenic EJ events by Sanger sequencing of the PCR product across the cut site of individual F2 flies, and subsequent analysis of the repair junction. MMEJ typically results from the annealing of direct repeats positioned on each side of the cut sites after resection, and deletion of the intervening sequence (Fig. 3A) [34, 66]. SD-MMEJ requires the presence of two ‘repeat motifs’, each containing a ‘primer’ (P) and a nearby ‘microhomology’ (MH) sequence [42] (Fig. 3A). Duplex unwinding or resection exposes the primer repeats that anneal to form loops or transient hairpins [42]. Primer annealing can occur between two direct repeats (‘loop-out’ mechanism) or two inverted repeats (‘snap-back’ mechanism) [42]. The 3′ end of the DNA in these structures then uses ssDNA as a template for DNA synthesis and repair. Unwinding of the secondary structure and re-annealing of the nascent DNA with a MH sequence on the other side of the break enables repair completion, typically resulting in deletions or indels (Fig. 3A).

**Figure 3:**
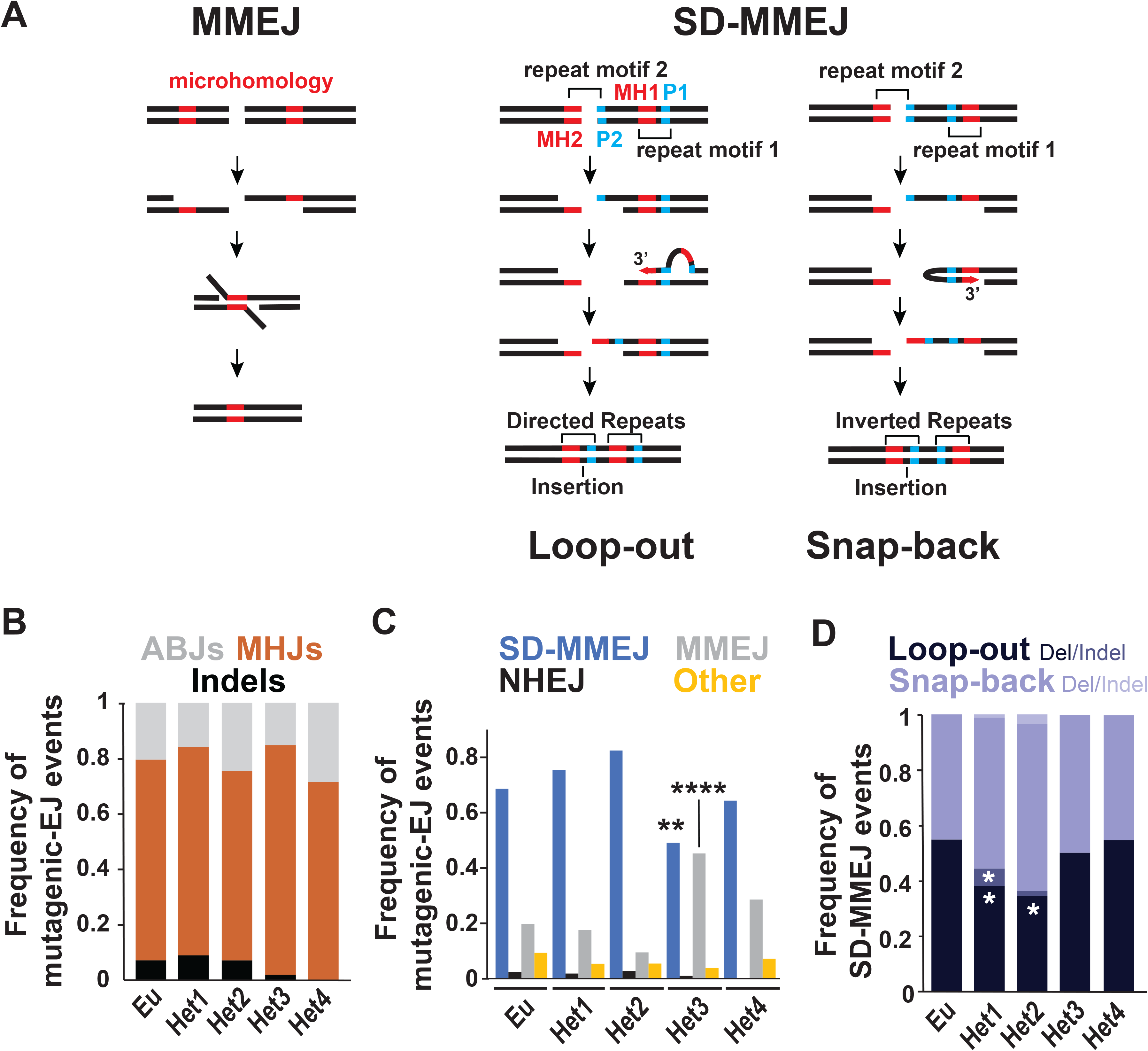
Frequency of alt-EJ repair pathways. A) Schematic representation of MMEJ and the main SD-MMEJ mechanisms (adapted from [16]), indicating the position of repeat motifs and the corresponding primers (P1, P2) and microhomologies (MH1, MH2). For the ‘loop-out’ mechanism, the DNA is unwinded before loop formation. For the ‘snap-back’ mechanism, resection occurs before hairpin formation. In both mechanisms, annealing of the break-proximal primer (P2) to the break-distal primer (P1), initiates DNA synthesis (red arrows) that creates new microhomology sequences (MH1) and that can create insertions. P1 and P2 are direct repeats for loop-out and inverted repeats for snap-back. Repair continues through secondary structure unwinding, annealing of the newly-synthesized microhomologies with MH2 sequences on the opposite side of the break, fill-in DNA synthesis, and ligation. When P2 and MH2 are not adjacent to the break site DNA flaps form and get trimmed, resulting in long deletions. When P1 and MH1 are adjacent to each other, insertions do not occur. B) Frequency of indels, MHJ and ABJ products of mutagenic-EJ repair across the different DR-*white* insertions. Sequences analyzed: n=86 for Eu; n=280 for Het1; n=74 for Het2; n=104 for Het3; n=14 for Het4. All comparisons are not significant. C) Frequency of SD-MMEJ-consistent, MMEJ-consistent (*i.e.,* SD-MMEJ-inconsistent MHJ), NHEJ (i.e, SD-MMEJ-inconsistent ABJ with less than 4bp-long alterations of the repair junction), or ‘other’ repair outcomes across different DR-*white* insertions. The ‘other’ category includes point mutations as well as ABJ and SD-MMEJ-inconsistent deletions or indels greater than 4bp. **p<0.005; ****p<0.0005 by two-proportion z-test, n=86 for Eu; n=280 for Het1; n=74 for Het2; n=104 for Het3; n=14 for Het4. D) Frequency of loop-out and snap-back-consistent SD-MMEJ outcomes across different DR-*white* insertions. n=59 for Eu; n=211 for Het1; n=61 for Het2. n=52 for Het3; n=9 for Het4; *p<0.05 by two-proportion z-test for the comparison between loop-out-consistent events of indicated heterochromatic DR-*white* insertions relative to the euchromatic one. All other comparisons are not significant.

Analysis of repair products across different DR-*white* insertions revealed that repair junctions are mostly characterized by deletions of less than 23 bp (Suppl. Fig. 2). The mean deletion size for all five insertions is between 9 and 16 bp (Suppl. Fig. 2), consistent with frequent repair by polθ-mediated alt-EJ mechanisms [34-36].

To establish the frequency of different mutagenic EJ events, we used previously published scripts in Python and R that identify repeat motifs and primer and microhomology repeats likely used during SD-MMEJ [42]. We classified repair products based on the presence of deletions or indels, which revealed that indels are rare for all DR-*white* insertions (less than 10% of all mutagenic EJ events) (Fig. 3B). This is consistent with previous studies that used DR-*white* as a repair reporter [21, 52]. We further classified repair events as deletions with apparent blunt joins (ABJs) and deletions with microhomology joins (MHJs) [37, 42]. This showed that deletions with microhomologies are the most common (∼70% of all mutagenic EJ events) across all DR-*white* insertions, in agreement with frequent MMEJ or SD-MMEJ repair (Fig. 3B).

Next, we established whether each repair event was consistent with SD-MMEJ mechanisms (Fig. 3C). To be classified as SD-MMEJ-consistent, the repair junction needs to satisfy the following requirements: i) the repeat motif needs to be at least 4 bp; ii) the primer repeat and the microhomology repeat within the repeat motif need to be at least 1 bp; and iii) the break-distal end of the repeat motif has to occur within 30 bp of the DSB [42]. MHJ repair events that are SD-MMEJ-inconsistent were identified as MMEJ. ABJ repair events or indels that are SD-MMEJ-inconsistent and result in a 1-4 bp deletion were classified as NHEJ [42]. ‘Other’ outcomes are rare and typically correspond to ABJ events associated with longer (5 bp or more) deletions. Notably, while we cannot exclude a contribution of cNHEJ to small (1-4 bp) deletions classified as SD-MMEJ consistent by the above criteria [33, 67, 68], previous studies in *Drosophila* showed that most SD-MMEJ-consistent deletions around the I-SceI cut site are Polθ-dependent and not Lig4-dependent, indicating a prevalent role of alt-EJ repair in this context [44].

This analysis reveals significant differences in the frequencies of usage of distinct mutagenic EJ pathways across the DR-*white* insertions (Fig. 3C). Specifically, Het3 is characterized by less SD-MMEJ-consistent and more MMEJ-consistent events relative to the euchromatic DR-*white* insertion (Fig. 3C). Within SD-MMEJ, loop-out deletions are significantly reduced in Het1 and Het2 relative to the euchromatic site, and loop-out indels are higher in Het1 (Fig. 3D). We conclude that there is variation in the frequency of use of distinct mutagenic EJ pathways across different DR-*white* insertions, without prevalence of a specific pathway in heterochromatin relative to euchromatin.

### SD-MMEJ repair results in smaller deletions in heterochromatin

Interestingly, the analysis of repair deletion size revealed differences between euchromatic and heterochromatic DR-*white* insertions (Suppl. Fig. 2). Het2 and Het3 are characterized by a significantly lower deletion size relative to the euchromatic site. Het1 and Het4 also show a lower mean and median size of deletions relative to the euchromatic site (Supplementary Fig. 2), although the difference in size distribution compared to the euchromatic site does not reach significance.

To understand these differences in greater detail, we first analyzed the position of the deletion boundaries associated with different repair mechanisms across all DR-*white* insertions (Fig. 4). This analysis reveals a higher frequency of boundaries close to the cut site for all heterochromatic insertions relative to the euchromatic one, resulting from SD-MMEJ- and/or MMEJ-consistent events (Fig. 4).

**Figure 4:**
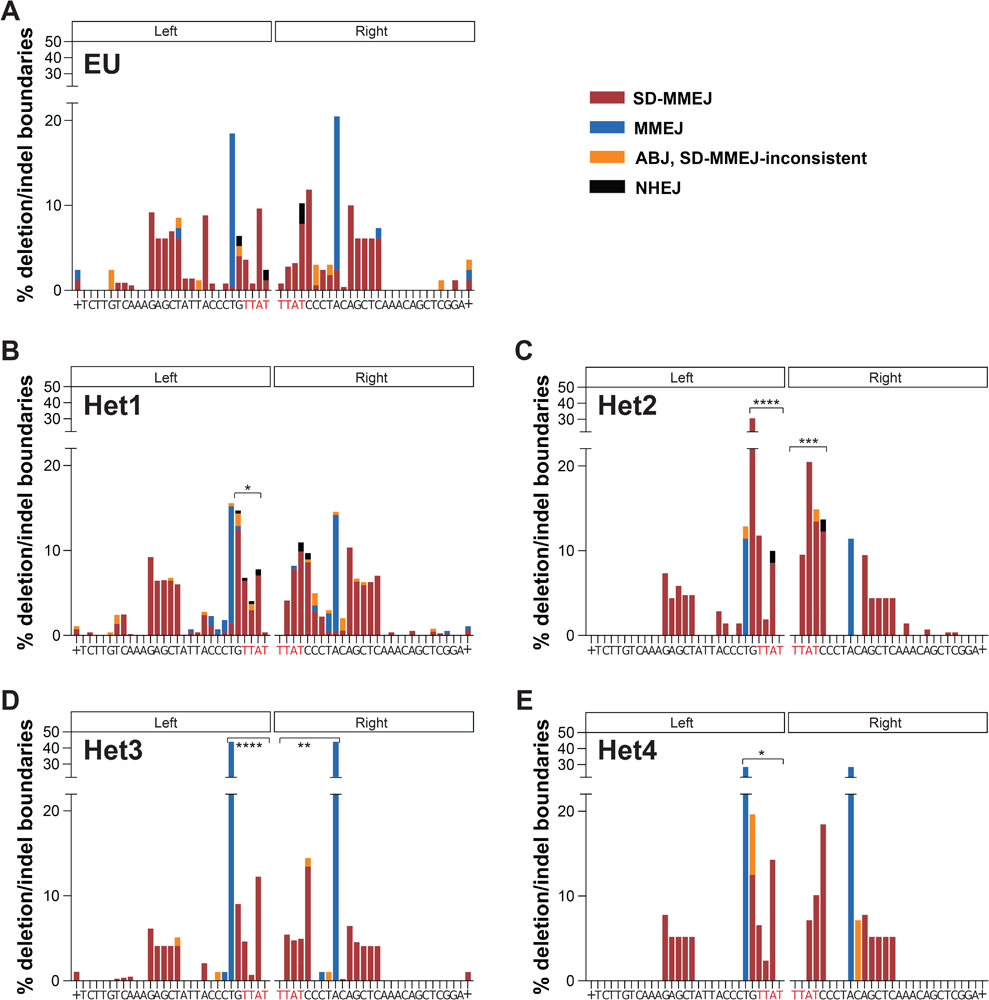
Distribution of deletion boundaries. Deletion boundaries associated with different alt-EJ pathways for Eu (A), Het1 (B), Het2 (C), Het3 (D), Het4 (E) DR-*white* insertions are shown. Deletion boundaries were defined as the first base proximal and first base distal to the deleted bases, with ambiguous bases aligned to the left of the cut site. + indicates all deletion boundaries to the left or right of the sequence shown. *p<0.05; **p<0.01; ***p<0.0005; ****p<0.0001 by two-proportion z-test for comparisons with the euchromatic DR-*white* insertion. n=86 for Eu; n=280 for Het1; n=74 for Het2; n=104 for Het3; n=14 for Het4 total repair events. Red text highlights the TTAT/AATA overhangs produced by I-SceI cutting.

In agreement with the analysis of deletion boundaries, a heat map plot of the frequency of deletion spreads for SD-MMEJ-consistent events highlights a prevalence of smaller deletions for all heterochromatic DR-*white* insertions relative to the euchromatic one (Fig. 5A). Conversely, the spread of deletion for MMEJ-consistent events remains largely the same across the different DR-*white* insertions, corresponding to repair using the CCCT/CCCT direct repeat proximal to the cut site (Fig. 4 and 5B). This indicates that, within SD-MMEJ-consistent events all heterochromatic DR-*white* insertions result in smaller deletions in heterochromatin relative to euchromatin, while deletions resulting from MMEJ-consistent events typically remain of the same size in the two domains.

**Figure 5:**
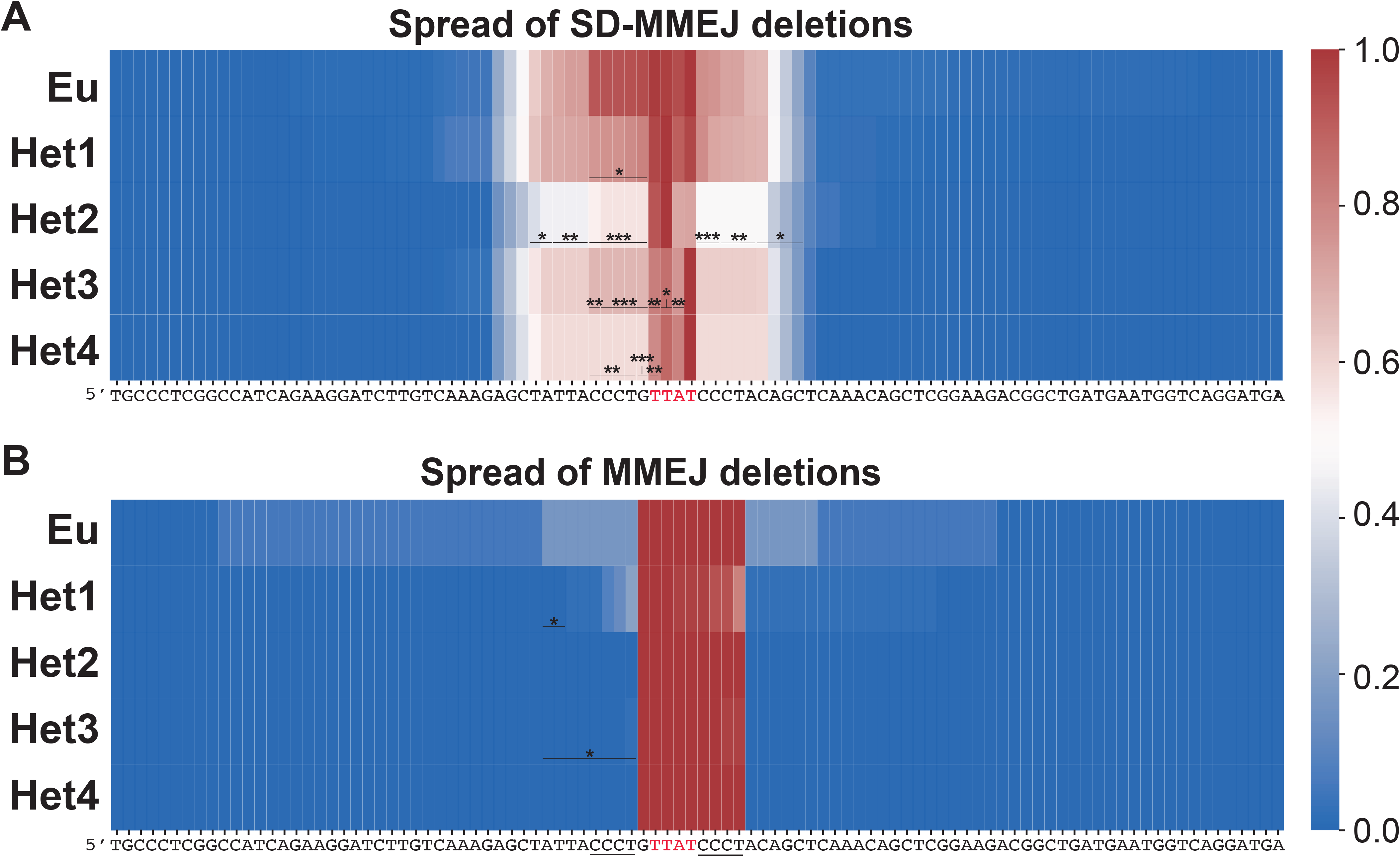
Spread of deletions for Mutagenic EJ repair. Heat map plots show the spread of deletions for SD-MMEJ-consistent (A) and MMEJ-consistent (B) repair events across the different DR-*white* insertions. The frequency of deletion occurring at each bp was calculated as the number of events with that bp deleted divided by the total number of events. Red text highlights the TTAT/AATA overhangs produced by I-SceI cutting. *p<0.05; **p<0.005; ***p<0.001 by two-proportion z-test at each bp for comparisons with the euchromatic DR-*white* insertion. (A) n=58 for Eu; n=198 for Het1; n=58 for Het2; n=52 for Het3; n=9 for Het4 total SD-MMEJ-consistent repair events. (B) n=19 for Eu; n=56 for Het1; n=7 for Het2; n=48 for Het3; n=5 for Het4 total MMEJ-consistent repair events.

Smaller deletions from SD-MMEJ repair could derive from the usage of repeat motifs positioned closer to the cut site in heterochromatin. We analyzed this directly by comparing the frequency of use of repeat motifs associated with different SD-MMEJ repair events (Fig. 6 and Suppl. Fig. 4). This analysis shows that the smaller deletion sizes for SD-MMEJ events in heterochromatin are typically associated with the use of repair motifs closer to the cut site for all heterochromatic DR-*white* insertions relative to the euchromatic site, particularly for repeat motif 2 (Fig. 6A,B).

**Figure 6:**
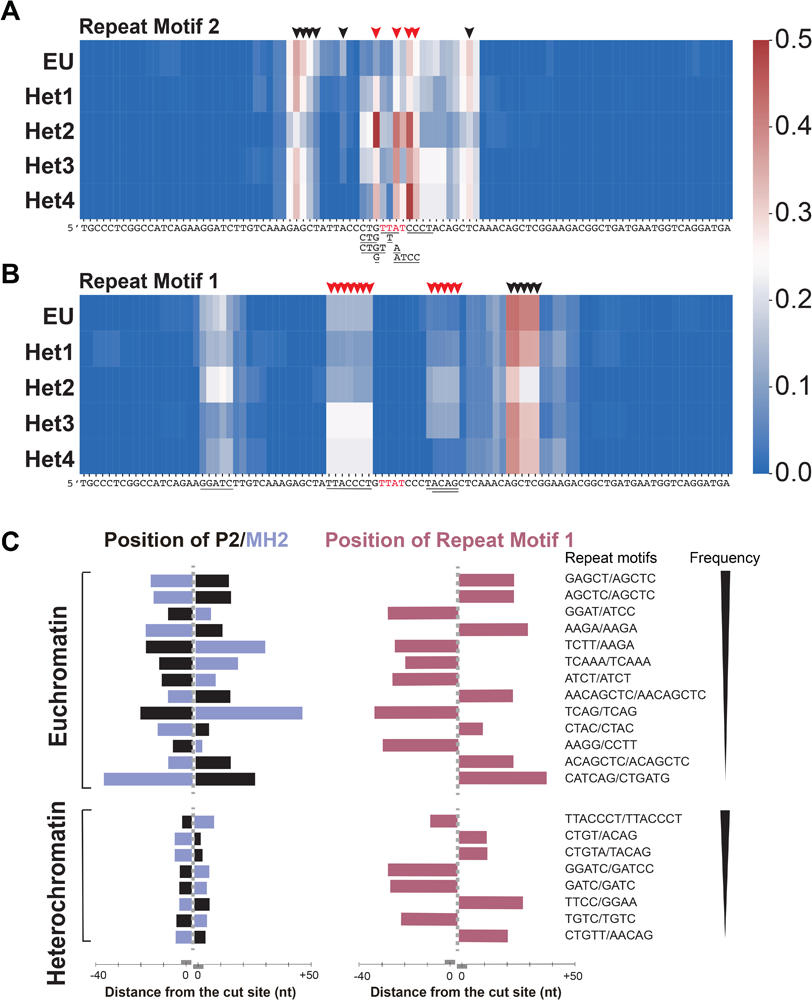
Position of repeat motifs for SD-MMEJ. Heat map plots show the frequency of usage of different repeat motifs 2 (A) and 1 (B) in SD-MMEJ repair resulting in deletions, for each DR-*white* insertion. Arrows highlight motifs typically used more frequently (red arrows) or less frequently (black arrows) in heterochromatic DR-*white* insertions, relative to the euchromatic one. n=58 for Eu; n=198 for Het1; n=58 Het2; n=52 for Het3; n=9 for Het4. Red text highlights the TTAT/AATA overhangs produced by I-SceI cutting. Underlined text highlights some of the most frequently used repeat motifs. (C) Schematic representation of the distance of P2/MH2 (left) and repeat motif 1 (right) from the cut site for the most common repair events in euchromatin and heterochromatin, sorted by frequency. The most common repair products are shown at the top of the list.

**Figure 7:**
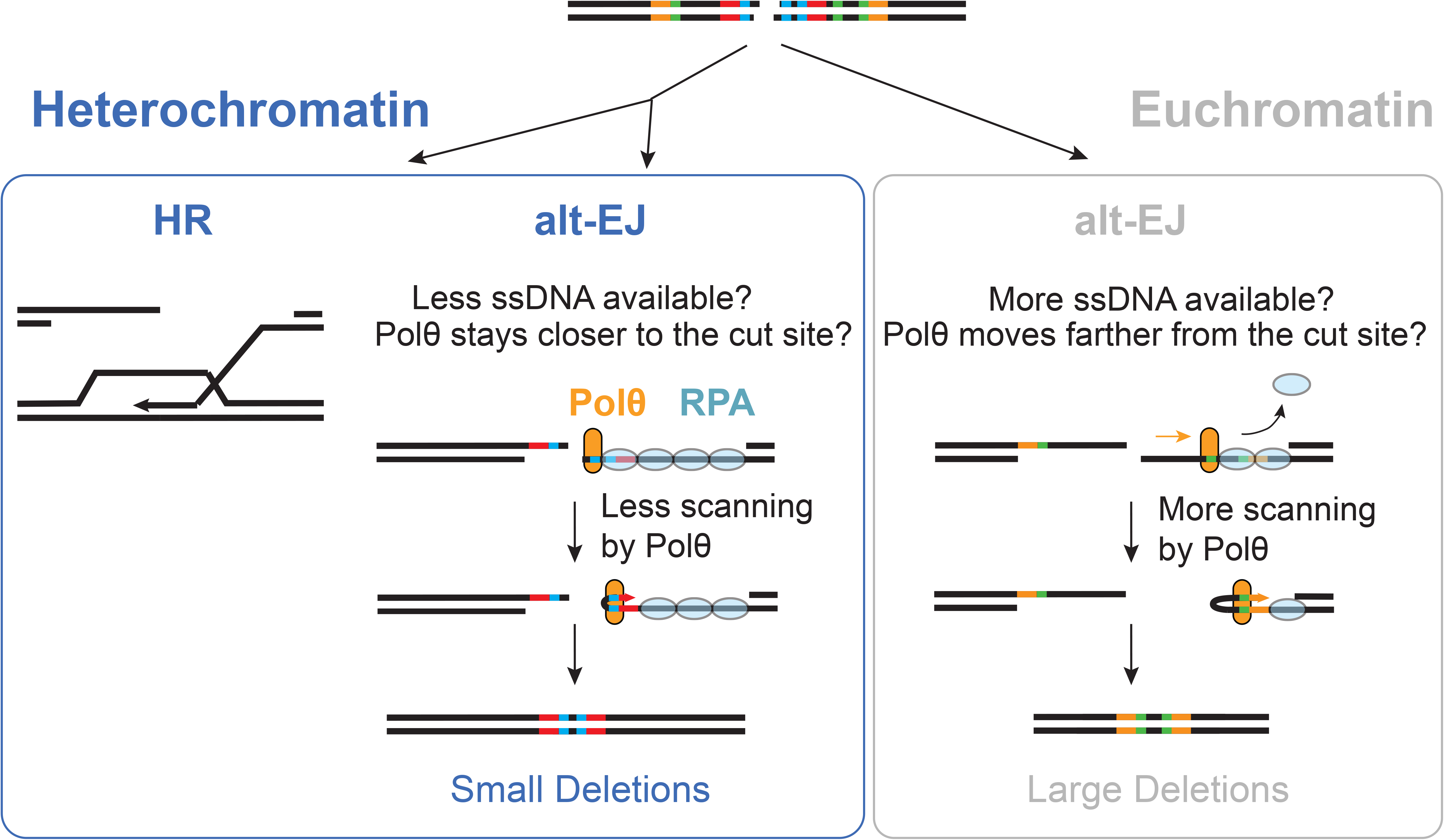
Model for HR and alt-EJ repair in heterochromatin. In heterochromatin (A) HR repair is promoted, and polθ binding close to the cut site facilitates the use of proximal primers for HR repair, resulting in smaller deletions. In euchromatin (B) polθ binding further from the cut site and more extensive probing for homology allows larger deletions to occur.

To further understand these differences, we identified the most frequent changes in repeat motifs usage between euchromatin and heterochromatin, corresponding to ∼70% of all the events (Fig. 6c and Suppl. Fig. 3). Mapping repeat motifs and primers used in these events confirmed a clear tendency for heterochromatic DR-*white* insertions to use a primer P2 and a microhomology MH2 closer to the cut site relative to the euchromatic one (Fig. 6c). The most common events in heterochromatin also rely on a Repeat Motif 1 located closer to the cut site than euchromatic events, although less common repair outcomes also use farther motifs. This suggests that polθ ability to probe for homology is also affected in heterochromatin, being constrained to shorter distances from the cut site, although polθ can occasionally find a Repeat Motif 1 at distances comparable with those found in euchromatin (Fig. 6c). Together, these results suggest that the initial association of polθ with the DNA end, and to some extent also homology probing and annealing for MH1 synthesis, tend to be confined to the region proximal to the cut site in heterochromatin. In addition, polθ typically finds homology (MH2) closer to the cut site in heterochromatin relative to euchromatin.

Together, we conclude that alt-EJ repair relies on primers and microhomologies closer to the cut site to initiate and complete alt-EJ repair in heterochromatin, resulting in SD-MMEJ repair with smaller deletions.

## DISCUSSION

Previous studies identified a clear role of the DNA sequence in affecting alt-EJ repair, but the role of the chromatin state in influencing alt-EJ remains poorly explored. Recent studies showed that lamin-associated silenced chromatin and H3K27me3-enriched chromatin interfere with cNHEJ while enabling MMEJ. However, no previous study directly addressed how alt-EJ repair operates in pericentromeric heterochromatin, a large region of the genome mostly composed of repeated DNA sequences, subject to epigenetic silencing by H3K9me2/3, characterized by phase separation, and mostly not associated with the nuclear periphery. There was also no clear understanding of whether chromatin states affect SD-MMEJ repair. To address these questions, here we use the DR-*white* repair reporter inserted in a euchromatic locus and several new locations in pericentromeric heterochromatin and investigate DSB repair responses in these contexts. We established an experimental system to induce damage in male spermatogonia, resulting in DSBs in a uniform population of actively dividing mitotic cells. With this setup, DNA sequence and cell cycle phases are the same for all the DR-*white* insertions analyzed, removing these factors as potential sources of variability in repair pathways. Our studies revealed a higher frequency of HR usage in heterochromatin relative to euchromatin. In addition, we identified a higher frequency of usage of microhomologies close to the cut site for SD-MMEJ repair in heterochromatin, resulting in smaller deletions. Together, these studies are consistent with the model that DSB repair is less mutagenic in heterochromatin than in euchromatin.

Detecting a higher frequency of HR repair in heterochromatin relative to euchromatin is in agreement with previous studies in *Drosophila* and human cells that suggested a preferential usage of HR for heterochromatin repair, when both HR and NHEJ are available [9, 22]. However, this result is not consistent with other studies in flies using the DR-*white* reporter, where a similar frequency of HR was detected between euchromatic and heterochromatic insertions of the cassette [21]. A possible reason for this discrepancy lies in the amount of silencing present in the DR-*white* cassette across different insertions used in the two studies. All DR-*white* insertions used for the study described here were exceptionally silenced, as demonstrated by the fact that HR products (*i.e.,* red-eyes) could only be revealed by crossing the F1 progeny with *su(var)205+/-* flies. This was not the case for heterochromatic DR-*white* insertions used in the previous study [21], consistent with a higher level of expression of the cassette.

Additionally, the induction of DSBs in spermatogonia provides unique advantages to the study of HR repair. First, these cells are mitotically dividing, resulting in a high number of cells in S/G2 with active HR repair (Fig. 1B). In addition, having a uniform population of cells introduces less variability in the system, and the lack of ‘jackpot’ events further limits noise. In agreement, we found more consistency in the frequency of HR outcomes across different experiments relative to what was previously shown with a Ub promoter-driven I-SceI in the germline [21], which likely facilitates the identification of differences across distinct DR-*white* insertions.

What mechanisms promote HR repair in heterochromatin still needs to be defined. An interesting possibility is that high levels of HP1 promote resection in heterochromatin, similar to the role of transient HP1 deposition previously described in human euchromatin [69-73]. Consistent with this hypothesis, live imaging of cultured cells showed that ionizing-radiation (IR)-induced repair foci of the ssDNA/RPA-associated proteins ATRIP and TopBP1 form faster and appear brighter in heterochromatin relative to euchromatin [9].

Our analysis of mutagenic-EJ repair revealed a high frequency of SD-MMEJ-consistent events, in agreement with previous studies with the DR-*white* reporter and I-SceI expression in the germline [52]. However, we detected a much broader set of SD-MMEJ and MMEJ-consistent repair junctions than in previous studies with the same reporter [52], suggesting that our experimental setup with *Bam*-dependent induction of I-SceI is particularly suitable for detecting microhomology-driven mutagenic EJ. This could be due to the frequent mitotic divisions occurring in spermatogonia, given that mutagenic EJ is a preferred pathway to repair DSBs remaining in the genome upon mitotic entry [74-76], and that I-SceI induction in these cells induces continuous cuts until a mutagenic (terminal) event occurs [52].

Interestingly, we detected significant variability in the type of alt-EJ repair pathway prevailing at each heterochromatic site, with MMEJ prevailing at Het3 and snap-back SD-MMEJ prevailing at Het1 and Het2. This suggests that factors other than the epigenetic composition, sequence composition, cell cycle phase, and association with the heterochromatin domain and its phase separated state, might be important to balance the frequency of SD-MMEJ and MMEJ mechanisms.

Alternatively, transient epigenetic changes and/or transcription in spermatogonia might affect this balance. For example, Het1 is located in a lncRNAs, while Het4 is inserted in a natural transposable element (TE) (Fig. 1D and Suppl. Table 1), and low levels of transcripts have been detected at these locations in testis [63] (Suppl. Fig. 4). Similarly, Het2 is associated with low levels of miRNAs [63] (Suppl. Fig. 4). Thus, an interesting possibility is that temporary or partial opening of the DR-*white* insertions occurs at these sites, influencing repair outcomes. In this light, the completely silenced state of Het3 [62, 63] (Fig. 1D and Suppl. Fig 5) might be the reason why MMEJ is particularly efficient at this site, suggesting MMEJ as a predominant mutagenic-EJ pathway operating in highly repeated and highly silenced heterochromatin.

Despite these differences, we also uncovered a general preference for the use of repeat motifs closer to the cut site in heterochromatin relative to euchromatin, resulting in smaller deletions. The reasons for this distinctive behavior remains unknown. It is tempting to speculate that features linked to a more compact chromatin or the phase separated state of the heterochromatin domain [15, 49, 50, 77-79] affect end-processing and/or repair protein accessibility to facilitate the use of microhomologies close to the cut site for SD-MMEJ repair.

Polθ uses a scanning mechanism to identify microhomologies near DSB ends, binding to and moving bidirectionally from the 3’ end [33, 80]. Our data suggest that the initial binding of Polθ to the ssDNA is particularly affected in heterochromatin, resulting in the use of a primer sequence P2 closer to the cut site relative to euchromatin. The Polθ scanning process seems to be also affected although to a lesser extent, with identification of homologies (Repeat Motif 2) typically close to the cut site for the most frequent repair outcomes.

A possible explanation for the association of Polθ closer to the cut site is the availability of shorter stretches of ‘naked’ ssDNA, *e.g.,* as a result of less resection or less/weaker RPA association nearby the DSB [46, 81, 82]. We tend to favor the second hypothesis, given that heterochromatic DSBs also experience more frequent HR relative to euchromatin (Fig. 2), which relies on resection. In addition, resection typically initiates through the formation of a nick positioned ∼30 bp from the cut site, generated by the Mre11 endonuclease, and which is followed by Mre11-dependent 3’-5’ exonuclease activity to resect the DNA back toward the break [83, 84]. The amount of ssDNA generated through this ‘short-range’ resection is sufficient to explain most alt-EJ repair events detected in both euchromatin and heterochromatin, consistent with resection not playing a major role in the observed differences.

On the other hand, RPA-mediated coating of the ssDNA could differ between euchromatin and heterochromatin, resulting in shorter tracts of ssDNA available for the initial Polθ binding. RPA has also been shown to bind ssDNA in distinct conformations [85, 86] (reviewed in [87]), resulting in different coverage of the ssDNA and distinct strength of association with the DNA. Polθ is able to remove RPA to enable the scanning process for alt-EJ [46, 47]. Thus, in heterochromatin shorter tracts of naked DNA at the cut site, looser association of RPA close to the cut site, or tighter RPA association further away from the cut site, might constrain the initial binding of Polθ to the region nearby the DSB and result in the use of more proximal primer P2 sequences. RPA association in a more ‘loose’ conformation or longer ssDNA tracts could also facilitate Polθ-mediated displacement of RPA for secondary structure formation, homology identification and annealing, resulting in more proximal repeat motif 1 usage. Intriguingly, RPA positioning is also potentially influenced by the position of the initial nick generated during resection. Specifically, studies in yeast showed that the Mre11 nicking activity occurs at a more distal site in active regions relative to silenced regions [84], which could affect RPA cooperative binding along the ssDNA and result in shorter tracts of ‘naked’ ssDNA close to the cut site in heterochromatin. In addition, the ATPase activity of Polθ is necessary for larger homology scanning [27], thus suppression of this activity could also account for less homology search in heterochromatin.

In this view, RPA association or Polθ activity could also be influenced by the selective permeability properties of the phase separated heterochromatin domain [15], and regulated by post-translational modifications which are also subject to selective filtering in phase separated structures [88].

Finally, an alternative mode of resection through Mre11 exonuclease activity has been described in human G1 cells [89], which generates ssDNA from the DSB end. Should this mode of resection be available in the experimental system and genomic locations described here, the compact chromatin state could delay resection in heterochromatin [90-92], favoring the use of repair motifs closer to the cut site. In agreement, nucleosomes have been shown to interfere with resection [93] and the more compact chromatin could exacerbate this barrier in heterochromatin.

Regardless of the exact mechanism, our study highlights the preferential use of HR or of alt-EJ pathways relying on microhomologies close to the cut site in heterochromatin, both of which reduce the mutagenic potential of repair mechanisms in heterochromatin. This could be advantageous, because about half of fly heterochromatin is composed of repeated DNA sequences where mutagenic events can quickly spread across satellites through recombinational processes that result in homogenization of the repeats, and which affect large genomic domains [94]. In addition, given the short size of the repeated unit for most *Drosophila* satellites [94], alt-EJ mechanisms using sequences close to the cut site could be particularly effective in this domain.

In sum, this study generated new experimental approaches to study repair responses to site-specific DSBs, and particularly alt-EJ repair, which are broadly applicable in the field. It also provides the first systematic characterization of alt-EJ pathways in euchromatin and heterochromatin, identifying distinctive outcomes in heterochromatin. Future studies are needed to establish which properties of the heterochromatin domain are responsible for these differences, how repair mechanisms are affected, how manipulations of the chromatin state can be applied to control repair mechanisms in gene editing applications, and how disease states (including cancer and aging) can affect these responses threatening genome stability.

## METHODS

### Drosophila stocks and maintenance

*Drosophila* stocks were maintained on standard media at 25°C, which was prepared as in [95]. We used the Minos-mediated integration cassette (MiMIC) system [96] to integrate the DR-*white* reporter [21] (DR-white-RFP) into four pericentromeric heterochromatic and one euchromatic loci (Supplementary Table 1). Embryo injection and validation of the insertions were performed by Bestgene Inc. (Chino Hills, CA) and Genetivision Corporation (Houstin, TX). BamP898-CFI (*Bam*-I-SceI) was described in [56]. All insertions were further verified by genotyping with locus-specific oligonucleotides (Table 1). Su(var)205^[1545]^ was described in [97] and was verified by sequencing across the mutation site.

### Crosses and experimental setup for the DR-*white* repair assay

Homozygous females containing DR-*white* on chromosome 2 were crossed to males containing BamP898-CFI on chromosome 3, for 3 days (F0) (Fig. 1F). Importantly, in BamP898-CFI the presence of the Bam 3’UTR enables tight suppression of protein synthesis in spermatocytes through post-transcriptional regulation [98]. This regulation is in addition to the transcriptional induction of Bam promoter that is limited to spermatogonia [54, 55, 56], resulting in a very tight I-SceI expression only in spermatogonia and not in spermatocytes. After enclosure, 0-1 day old males containing DR-*white* and *Bam*-I-SceI were individually crossed to Su(var)205^[1545^]/Cyo females (F1). The parents were removed after 3 days and the F2 progeny containing DR-*white* and Su(var)205^[1545]^ was collected for visual scoring and sequence analysis. We obtained between 21 and 47 vials with viable F1 crosses for each experiment, resulting in 200-926 DR-*white*-containing F2 flies that were used for the different analyses.

### DR-*white* assay

F2 flies with w+ phenotype (red eyes) were classified as HR, while flies with phenotype w-DsRed-were classified as SSA and confirmed by PCR, as previously described [21, 52]. Only about 40% of the flies scored as DsRed- was confirmed to contain a SSA product by PCR, the rest was identified as an EJ product after sequencing. This indicates that DsRed is not fully penetrant or easy to score in the F2 progeny with our experimental setup. w-DsRed+ flies were used for the molecular analysis of EJ-products as follows. Individual F2 males were processed for DNA extraction, PCR amplification, and I-SceI digestion, as previously described [52]. PCR products were obtained using Q5 Polymerase (NEB) or Expand Long Template PCR system (Roche/Sigma) and the oligonucleotides: DR-*white*2.2 and DR-*white*2.2a (Suppl. Table 1). Sanger sequencing of PCR products that lost the I-SceI cut site was performed by Laragen (Culver City, CA) using an oligo 420 bp from the I-SceI recognition site (Suppl. Table 1), after an additional purification step.

### Analysis of mutagenic EJ events

Sequences were trimmed to common start and end positions, and mapped to the reference DR-*white* sequence using Clustal Multiple Sequence Alignment tool. This alignment revealed sequences with insertions and deletions. Deletion junctions were analyzed using a previously published Python script [42], classified as apparent blunt joins (ABJ) or microhomology junctions (MHJ), and further analyzed for SD-MMEJ consistency. Next, SD-MMEJ-consistent junctions were analyzed to identify repeat motifs of 4 or more bp in the repair product, and the corresponding loop-out (for direct repeat motifs) or snap-back (for inverted repeat motifs) mechanisms associated with their usage [42] (Fig. 3A and Suppl. Fig. 4 for examples of SNAP-back events). For deletion junctions characterized by several motif-deletion pairs that could result in the same repair outcome (examples are shown in Suppl. Fig. 6), each motif-deletion pair was assigned a weight assuming each pair is used with the same probability, as in [42]. Indel sequences (containing novel sequences not present in the original construct, with or without deletions) were analyzed using a previously described R script to determine SD-MMEJ consistency and identify motifs, primers, and microhomologies [42]. Notably, some insertions can be generated by several rounds of annealing, synthesis, and dissociation prior to the final pairing [33, 37], but the computational models applied here assume that all SD-MMEJ-consistent events are generated from a single templated synthesis and microhomology annealing event [42]. Multi-step events are likely responsible for some of the outcomes indicated as ‘other’ in Figure 3C.

### Quantification of H3K9me3 enrichments at insertion sites

ChIP-seq data were from Kc cells ([61] and this study). ChIP experiments were conducted as in [99] using Ab8898 antibodies (Abcam), and immunoprecipitated samples were processed for 150bp pair-end sequencing on Hi-Seq platform of Illumina by Novogene. For all replicates and input, sequencing reads were trimmed of their adapters and low quality reads were removed using trim galore. Reads were aligned to the *Dm6* reference genome using the BWA-MEM alignment algorithm. Optical duplicates were removed using MarkDuplicates (Picard). Coverage within +/- 0.5 Mb of each insertion site was determined using Rsamtools. H3K9me3 and input read counts within +/- 0.5 Mb of each insertion site were normalized for sequencing depth using RPM normalization. Enrichments calculated over other intervals between 1Kb and 100Kb gave similar results.

### Quantification of miRNA-seq and RNA-seq enrichments at insertion sites

*Drosophila* testis miRNA-seq data from [63] was obtained from the Sequence Read Archive with accession numbers: ERR2117087, ERR2117088, ERR2117089. *Drosophila* testis RNA-seq data from [63] was obtained from SRA with accession numbers: ERR2105061, ERR2105062, ERR2105063. Reads were trimmed of their adapters, low quality reads were removed using Trim Galore, and were aligned to the Dm6 reference genome using Bowtie. Read counts within +/- 10 kb each DR-*white* insertion site were calculated as reads per kilobase per million reads (RPKM; Suppl. Fig. 5)). Read counts for spermatogonia-specific reference RNAs were also calculated as RPKM.

## Acknowledgments

This research was supported by NIH R01GM117376 and NSF Career 1751197 to I.C, and MCB1716039 to M.M. We are grateful to Tzumin Lee for sharing BamP898-CFI flies before publication, and Gary Karpen for sharing Su(var)205^[1545]^ flies. We thank Cuiyi Zhang for helping with fly genotyping, Haseeb Khan for helping with PCRs of F1 flies, and Nicholas Woodward for critical reading of the manuscript. MiMIC stocks obtained from the Bloomington Drosophila Stock Center (NIH P40OD018537) were used in this study.

## Competing interests

The authors declare no competing interests.

**Supplementary Figure Legends:**

---

**Supplementary Fig. 1:**
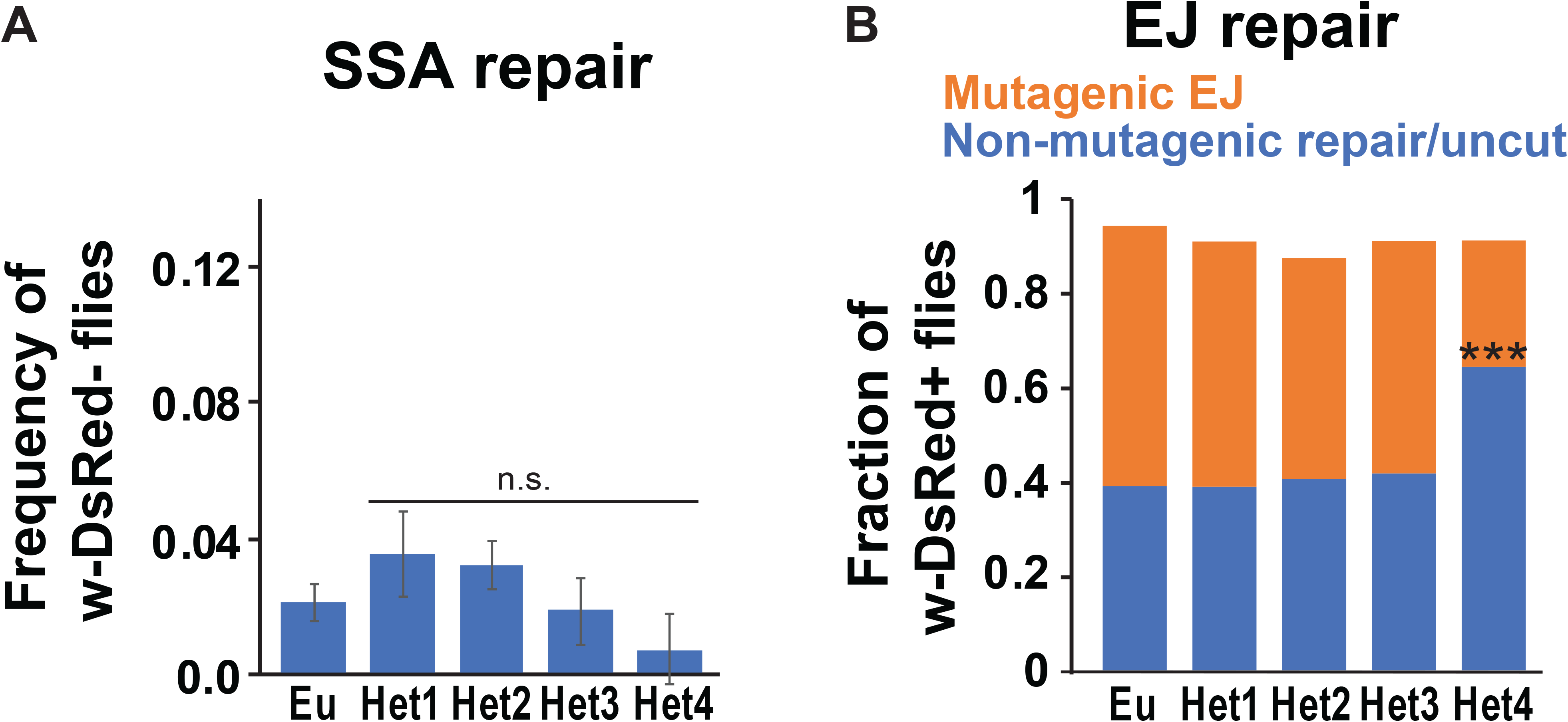
Mutagenic repair frequency. Frequency of A) SSA, or B) mutagenic-EJ and non-mutagenic repair/uncut, in white-eyed F2 flies is shown. ***p<0.001 by two proportion z-test. n.s.: not significant. Mutagenic repair results in a PCR product across the cut site that cannot be digested by I-SceI. Non-mutagenic repair results in a PCR product across the cut site that can be digested by I-SceI. In A) n=1352 for Eu; n=1267 for Het1; n=880 for Het2; n=679 for Het3; n=485 for Het4. Error bars: SEM. In B) n=570 for Eu; n=193 for Het1; n=166 for Het2; n=312 for Het3; n=85 for Het4.

**Supplementary Fig. 2:**
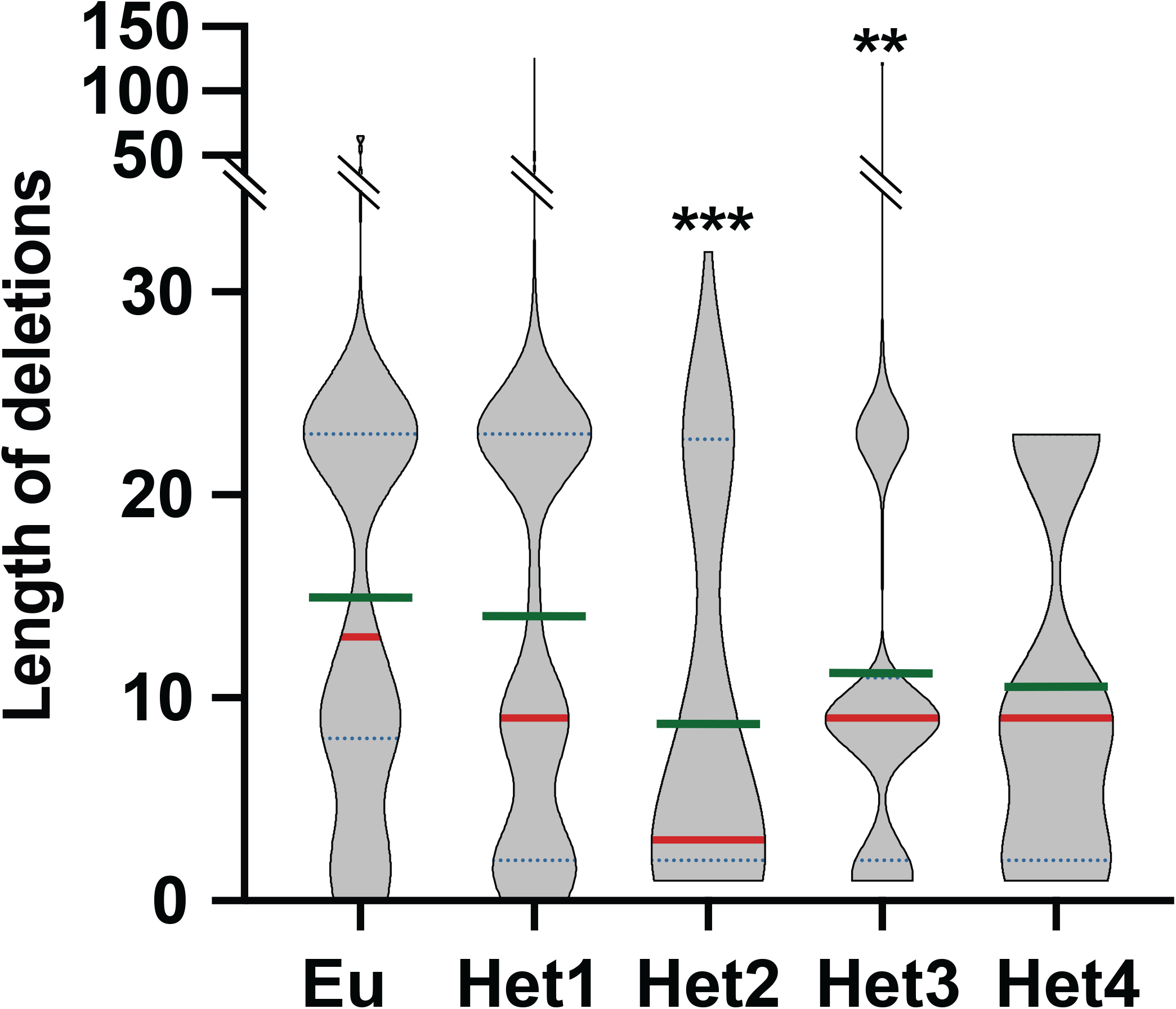
Average size of deletions. Violin plots show the distribution of deletion sizes across all DR-*white* insertions. Green line: mean. Red line: median. Dashed black lines: quartiles. **p<0.01; ***p=0.0003 by one-tailed Mann-Whitney test. n=86 for Eu; n=280 for Het1; n=72 for Het2; n=104 for Het3; n=14 for Het4.

**Supplementary Fig 3:**
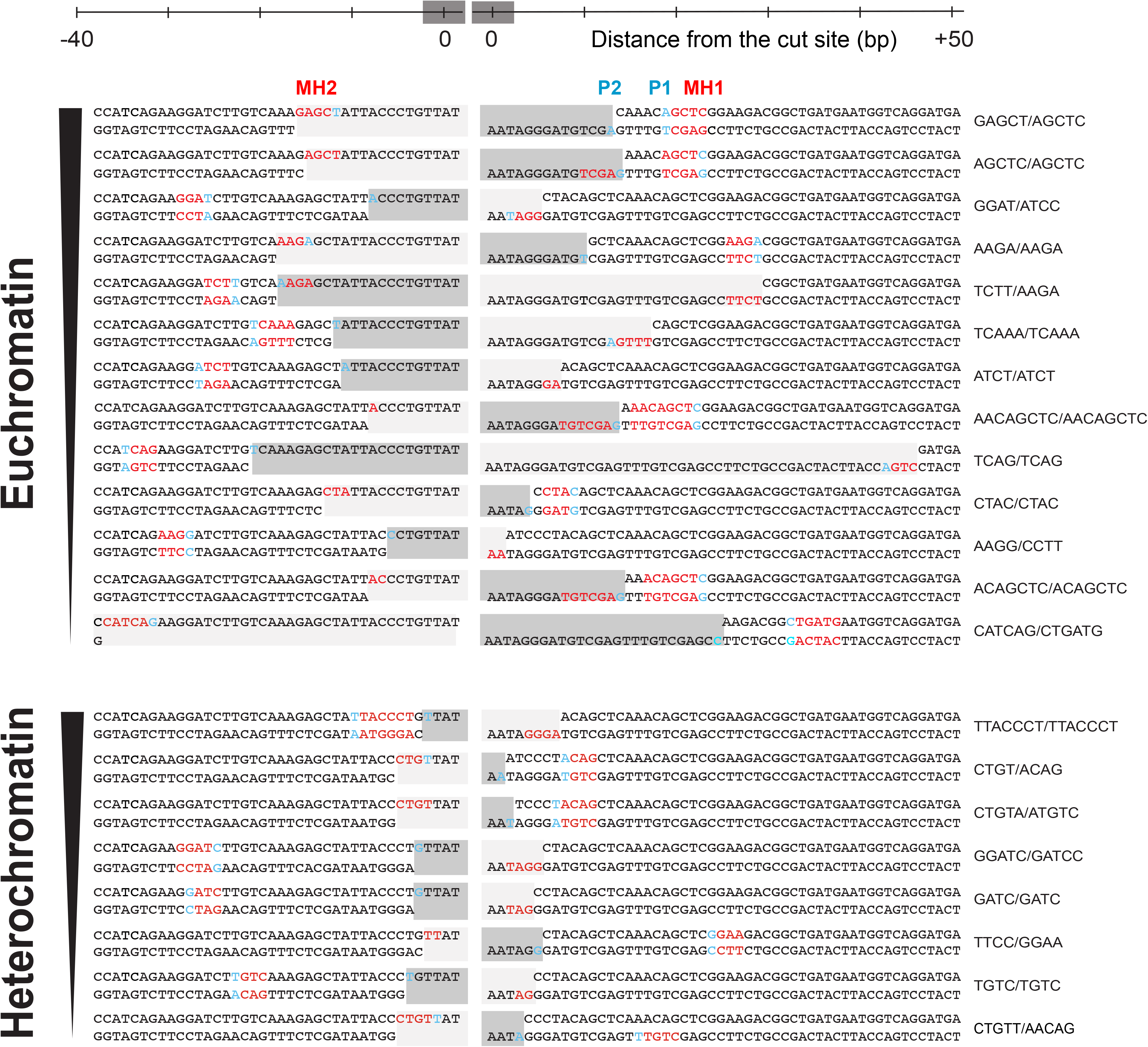
Common SD-MMEJ repair events in euchromatin and heterochromatin. The sequences of the most common SD-MMEJ repair events shown in Fig. 6C, have been sorted from top to bottom by frequency of occurrence. Red: microhomologies. Blue: primers. The minimum primer sequence is shown for all repair outcomes.

**Supplementary Fig. 4:**
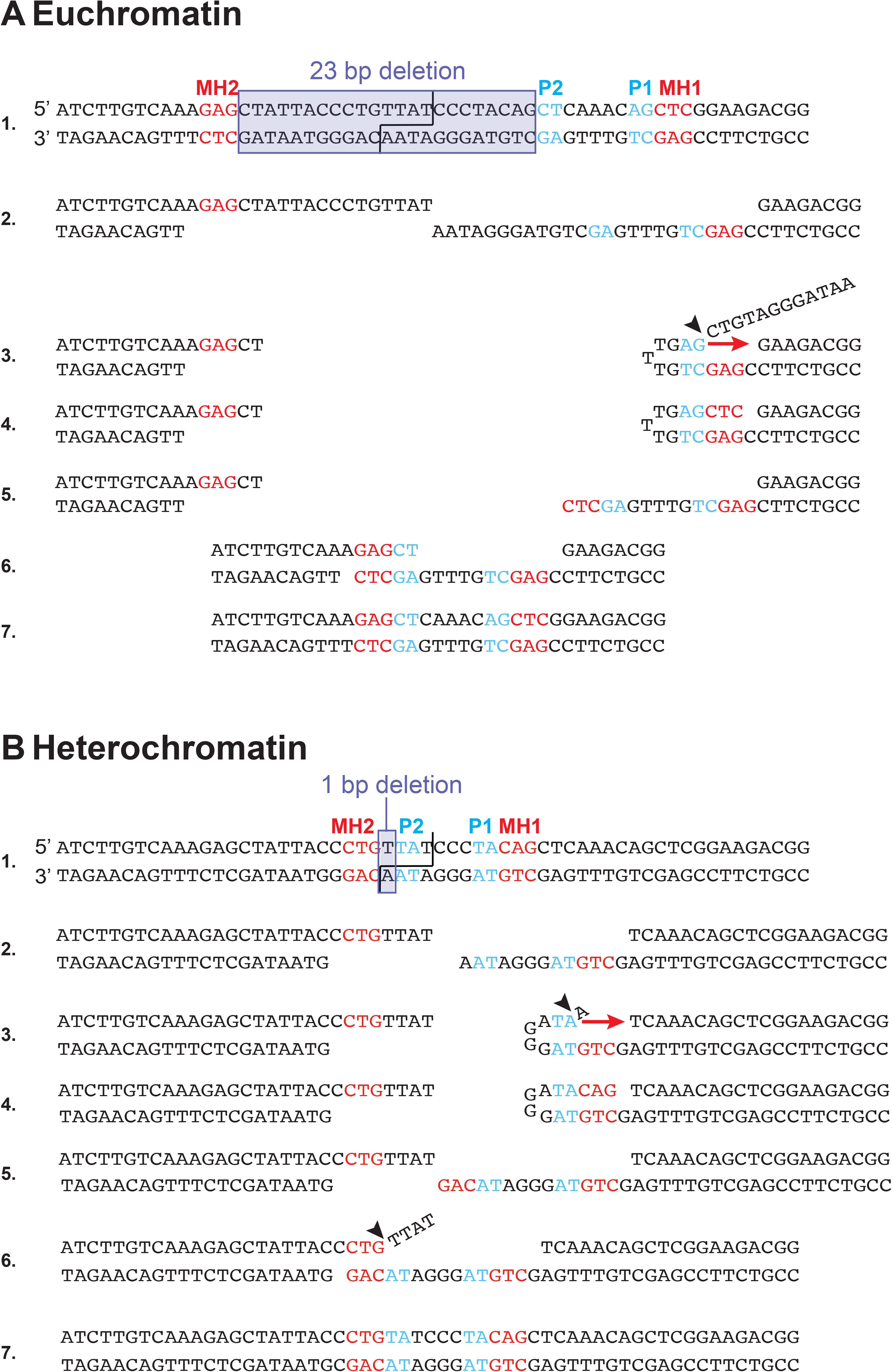
Common SD-MMEJ intermediates in euchromatin and heterochromatin. Examples of snap-back SD-MMEJ repair intermediates (1-7) for two of the most frequent repair events detected in euchromatic (A) and heterochromatic (B) DR-*white* insertions, highlighting: the deleted tract (violet square); the cut site (black line); the position of primers (P1, P2) and microhomologies (MH1, MH2) as defined in Figure 3A; templated DNA syntheses (red arrows) and flap removals (black arrowheads).

**Supplementary Fig. 5:**
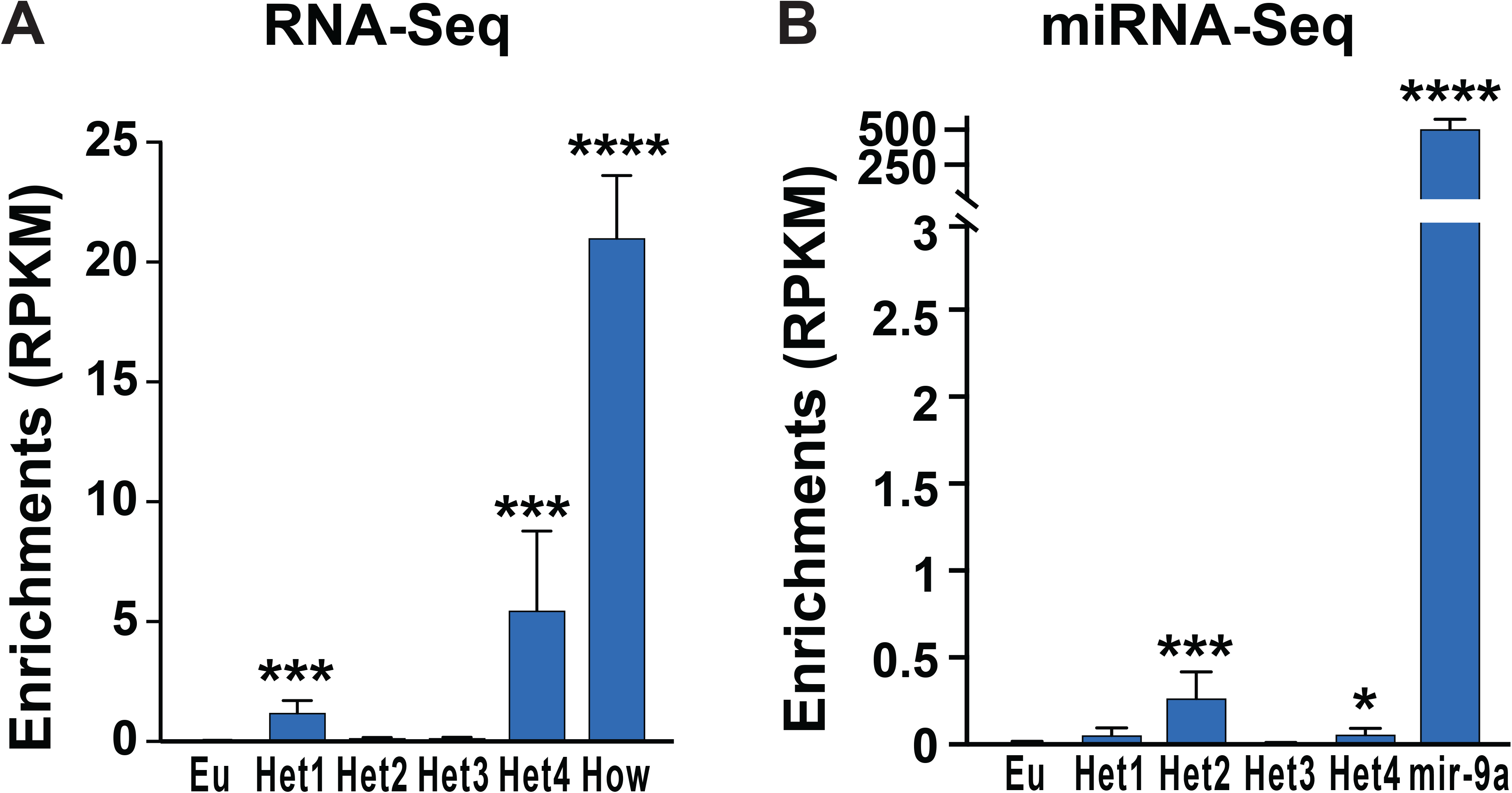
miRNA-seq and RNA-seq enrichments at euchromatic and heterochromatic insertion sites. Average reads per kilobase per million reads (RPKM) values of A) RNA-seq and B) miRNA-seq data from *Drosophila* testes at +/- 10Kb from DR-*white* insertion sites [63]. *How* and *mir-9a* are spermatogonia-enriched RNAs, used as a reference [100, 101]. ***p<0.005, ****p<0.0001 by two-tailed two-sample z-test for means. Error bars: SD of 3 replicates.

**Supplementary Fig. 6:**
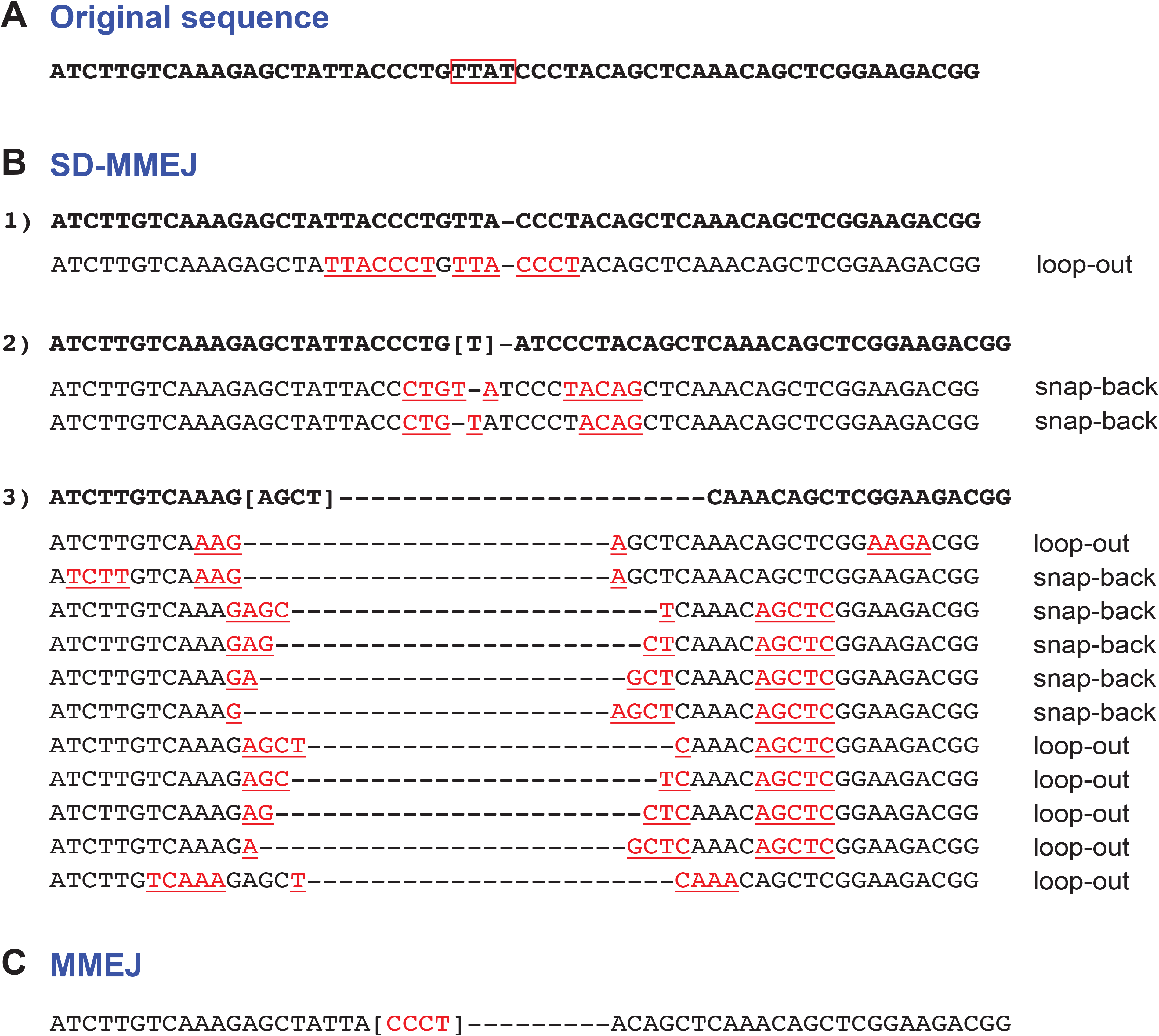
Common SD-MMEJ and MMEJ repair outcomes. The most frequent deletions across all DR-*white* insertions are shown (B,C), relative to the original sequence around the I-SceI site of the *SceI.white* (A). The red rectangle highlights the region containing overhangs from I-SceI cut. B) Three examples (1-3) of SD-MMEJ-consistent repair outcomes are shown, each followed by the possible deletions and repeat motifs (red, underlined) associated with it. Brackets mark bases that could align to the left or to the right of the break, and cannot be unequivocally assigned to one position. In the analysis, all possible outcomes are assigned a weight assuming equal probability of occurrence. The respective SD-MMEJ repair mechanism (snap-back or loop-out) is also indicated. C) The microhomology used for MMEJ repair is shown in brackets.

**Supplementary Table 1.**
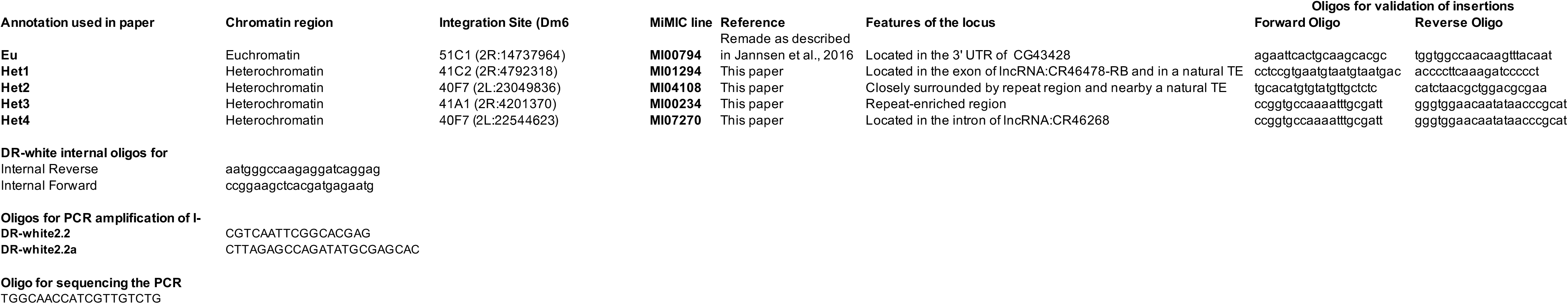

